# From STRs to SNPs via ddRAD-seq: geographic assignment of confiscated tortoises at reduced costs

**DOI:** 10.1101/2021.12.07.471568

**Authors:** Roberto Biello, Mauro Zampiglia, Silvia Fuselli, Giulia Fabbri, Roberta Bisconti, Andrea Chiocchio, Emiliano Trucchi, Daniele Canestrelli, Giorgio Bertorelle

**Author notes:** Corresponding authors Roberto Biello Giorgio Bertorelle.

## Abstract

Assigning individuals to their source populations is crucial for conservation research, especially for endangered species threatened by illegal trade and translocations. Genetic assignment can be achieved with different types of molecular markers, but technical advantages and cost saving are recently promoting the shift from short tandem repeats (STRs) to single nucleotide polymorphisms (SNPs). Here, we designed, developed, and tested a small panel of SNPs for cost-effective geographic assignment of individuals with unknown origin of the endangered Mediterranean tortoise *Testudo hermanni*. We started by performing a ddRAD-seq experiment on 70 wild individuals of *T. hermanni* from 38 locations. Results obtained using 3,182 SNPs are comparable to those previously obtained using STR markers in terms of genetic structure and power to identify the macro-area of origin. However, our SNPs revealed further insights into the substructure in Western populations, especially in Southern Italy. A small panel of highly informative SNPs was then selected and tested by genotyping 190 individuals using the KASP genotyping chemistry. All the samples from wild populations of known geographic origin were genetically re-assigned with high accuracy to the original population. This reduced SNPs panel represents an efficient molecular tool that enables individuals to be genotyped at low cost (less than €15 per sample) for geographical assignment and identification of hybrids. This information is crucial for the management in-situ of confiscated animals and their possible re-allocation in the wild. Our methodological pipeline can easily be extended to other species.

## 1 INTRODUCTION

In the last twenty years, Next-Generation Sequencing (NGS) favoured a shift from short tandem repeats (STRs) markers to single nucleotide polymorphisms (SNPs) in molecular studies of many organisms (Barbosa, Hendricks, Funk, Rajora, & Hohenlohe, 2020; Rajora, 2019; Seeb et al., 2011). This transition improved our understanding in several areas relevant to conservation, such as the study of inbreeding (Grossen, Guillaume, Keller, & Croll, 2020; Robinson et al., 2019), hybridization (Sinding et al., 2018; VonHoldt et al., 2016), population structure and admixture (Jeffries et al., 2016; Zimmerman, Aldridge, & Oyler-McCance, 2020), and population, parentage and kinship assignment (Kleinman-Ruiz et al., 2017; Roques, Chancerel, Boury, Pierre, & Acolas, 2019).

Some of the most common and widespread techniques used for genotyping SNPs in non-model organisms are based on reduced libraries as the restriction site-associated DNA sequencing (RAD-seq; Davey et al., 2011; Hohenlohe, Amish, Catchen, Allendorf, & Luikart, 2011; Miller, Dunham, Amores, Cresko, & Johnson, 2007), its variant double-digest RAD-seq (ddRAD-seq; Peterson, Weber, Kay, Fisher, & Hoekstra, 2012) and other derived methods (Campbell, Brunet, Dupuis, & Sperling, 2018). Currently, sequencing of nuclear genomic regions with reduced library approaches offers the possibility to obtain, at low price, several thousands of SNPs. Moreover, selecting *a posteriori* only the informative SNPs that maintain as much as possible the original dataset information (e.g., SNP-based panel) required for the purpose of interest can considerably reduce the subsequent genotyping costs and technical efforts necessary to analyze large number of individuals. This aspect is crucial in conservation genetics where practitioners need accuracy but also cheap and easy-to-implement guidance (Holderegger et al., 2019). SNP-based panels, compared to STRs, address this need by simplifying the allele scoring procedure, allowing data transferability across studies, with the potential for high-throughput screening (e.g., Garvin, Saitoh, & Gharrett, 2010). SNP arrays are also flexible, with low-density arrays allowing tens to thousands of SNPs to be genotyped and high-density arrays or “SNP chips” supporting tens of thousands to millions of SNPs (e.g., Humble, Paijmans, Forcada, & Hoffman, 2020; von Thaden et al., 2020).

Until recently, reduced SNP assays were used mostly for domestic species (e.g., Johnston, Huisman, Ellis, & Pemberton, 2017; Ogden, Baird, Senn, & McEwing, 2012; Pertoldi et al., 2009). Nowadays, custom species-specific SNP panels are being developed for several wild species providing insights into diverse topics including conservation genomics (e.g., Eriksson, Ruprecht, & Levi, 2020; Feutry et al., 2020; Henriques et al., 2018; Meek et al., 2016). Small panels of SNPs (<100) have already been used for the identification of distinct genetic populations to delineate biologically accurate management units and for the identification of individuals, including the assignment of individuals to their source population (Förster et al., 2018; Henriques et al., 2018; Kleinman-Ruiz et al., 2017).

This approach can be described in four steps. First, a large number of SNPs is isolated and typed in a small number of individuals; second, the suitability of these markers to answer the question of interest is tested; third, a much smaller panel of SNPs is identified, given it can answer the same question of interest but at lower costs; fourth, the subset is tested using a larger sample of individuals.

Here we applied these steps to isolate informative SNPs for the assignment of samples to their population or area of origin. Inferring the geographic origin of living organisms from their genotypes is of great interest in wildlife management, conservation, and forensic applications. It can provide information about gene flow, migration patterns and connectivity in natural populations (Kremer et al., 2012) but can also help inform wildlife managers about illegal animal translocations and poaching hotspots (Biello et al., 2021; Ogden & Linacre, 2015). The development of molecular tools for identifying the geographic origin of a specimen in a forensic context is still in its early stages (Ogden & Linacre, 2015). Our study produced a tool specific for the Herman’s tortoise, but it can be considered also as a useful example to facilitate the production of reduced SNP panels in other species.

The Hermann’s tortoise, *Testudo hermanni* Gmelin (1789), is one of the most endangered reptiles in Europe. The intense harvesting for the pet trade, especially before the 1980s when it was not banned yet (Ljubisavljević, Džukić, & Kalezić, 2011), the releases of non-native individuals, and the habitat reduction and degradation (Stubbs, Swingland, Hailey, & Pulford, 1985), are the major threats for this species (Bertolero, Cheylan, Hailey, Livoreil, et al., 2011). *Testudo hermanni* is distributed in disjoint populations across Mediterranean Europe, from Spain to the Balkans, including various Mediterranean islands. Based on mtDNA and STR markers, two subspecies are recognized (Biello et al., 2021; Perez et al., 2014): the eastern *T. h. boettgeri* and the western *T. h. hermanni*. The species is included in the list of strictly protected fauna species by the Bern Convention on the Conservation of European Wildlife and Natural Habitat. The western subspecies, *T. h. hermanni*, is classified as “Endangered” by the IUCN Red List (1996). Previous genetic studies suggest that individuals can be correctly assigned in most cases to their macro-area of origin with a small panel of STR markers (Biello et al., 2021; Perez et al., 2014). However, given the large number of individuals of unknown origin hosted in rescue centres and potentially suitable for reintroductions plans, the possibility to use modern NGS technologies to decrease the genotyping costs should be carefully considered.

The main goal of this study is to introduce and validate a small panel of SNPs extracted from genome-wide distributed markers for the cost-effective geographic assignment of large numbers of confiscated *Testudo hermanni* individuals, necessary for their management and their possible re-allocation in the wild. We accomplished this goal following the four steps defined above. In particular, 1. we performed a ddRAD-seq experiment on 70 wild individuals; 2. we verified that the results provided by the ddRAD-seq loci, in terms of genetic structure and power to identify the macro-area of origin, are adequate (in our case, considering the results obtained with a sample of 292 wild individuals already typed at 7 STRs); 3. we designed a cost-effective reduced panel of SNPs with retained informativeness; 4. we tested the reduced SNP panel in 190 individuals using the KASP-by-Design Fluidigm Assays (LGC Genomics) (He, Holme, & Anthony, 2014). Our study has practical consequences for the conservation, management, and reintroduction of *Testudo hermanni*, and it is also demonstrative of a technical pipeline applicable to other species.

## 2 MATERIAL & METHODS

### 2.1 Samples, DNA extraction and ddRAD-seq library preparation

Eighty-four samples already available in our laboratories were selected to perform the ddRAD-seq genotyping experiment. These samples belonged to individuals collected across most of the contemporary geographical range of the species and covering all the previously known genetic clusters.

DNeasy Blood and Tissue Kit (Qiagen) and Quick-DNA Universal Kit (Zymo Research) were used to extract genomic DNA from paper and whole blood stored at −20 °C, respectively, following manufacturer instructions. DNA was quantified with Qubit using the dsDNA BR kit (Invitrogen). ddRAD-seq libraries were prepared using *Sbf*I and *Nco*I for restriction digestion. Sequencing was performed on the Ion Torrent PGM with the Ion PGM Hi-Q View Sequencing kit (Life Technologies) following manufacturer’s instructions. See **Supplementary Methods** for more details about library preparation and sequencing.

### 2.2 Quality control and filtering

Raw reads generated were demultiplexed and split into individual data files using the *process_radtags* program in version 1.44 of STACKS (Catchen, Amores, Hohenlohe, Cresko, & Postlethwait, 2011). We used TRIMMOMATIC (Bolger, Lohse, & Usadel, 2014) to trim our raw reads, removing read-through adapters, low quality (light filter - modify on a case-by-case basis), and shorter than 200 bp, cropping the first two bp (GG left from restriction site). Resulting reads were 200 bp in length to ensure downstream compatibility in STACKS, which requires uniform read length when building *de novo* loci.

After trimming and quality-filtering, the STACKS programs *ustacks*, *cstacks* and *sstacks* were used to build de novo catalog loci. We specified a minimum of five reads per locus to build primary catalog stacks and allowed a maximum distance of six nucleotide mismatches within these stacks. Furthermore, we allowed eight mismatches between stacks during construction of the catalog because the specimens in our dataset were represented by two subspecies of the same species.

Following de novo mapping, an initial data-filtering step was performed using the *population* component of STACKS retaining only those loci present in at least 75% of individuals at each site and with a maximum of 5 SNPs per locus (*full dataset*). In an additional filtering step, we retained only one SNP per locus (*single SNP dataset*) to minimize redundancy of genetic information due to the physical linkage between the markers. This reduced dataset was used for PCA and F_ST_-based methods. The resulting vcf files were converted to other program-specific input files using PGDSPIDER v2.0.5.1 (Lischer & Excoffier, 2012).

### 2.3 Genetic structure with ddRAD-seq in comparison with STRs

In order to compare the results provided by the ddRAD-seq loci, in terms of genetic structure and power to identify the macro-area of origin of the individuals, we selected a subset of 292 wild samples genotyped previously at 7 STR loci (Biello et al., 2021; Perez et al., 2014) from the same locations of the samples included in the ddRAD-seq experiment.

The population structure was inferred from the ddRAD-seq with the package *fineRADstructure* designed to identify co-ancestry from RAD-seq data (Malinsky, Trucchi, Lawson, & Falush, 2018). Briefly, the algorithm compares each RAD-seq locus of each individual (recipient) with the alleles in all other individuals (potential donors) estimating the number of sequence differences (i.e., SNPs) to infer its nearest neighbor allele (donor) that is the allele with the least number of differences. The local co-ancestry values are then summed across all loci to obtain the co-ancestry similarity matrix for the full data. Next, an MCMC clustering algorithm infers the most likely population configuration. In summary, using information from the entire genetic variation identified with RAD-seq, the analysis estimates the number of genetic clusters, quantifies ancestry sources in each group in terms of co-ancestry value, and provides a tree-like illustration of the relationships between clusters. To perform this analysis, the output of the software *population* in STACKS was converted into a *RADpainter* input file with the python script Stacks2fineRAD.py. Samples were assigned to populations using 100,000 iterations as burn-in before sampling 100,000 iterations. A tree was built using 10,000 iterations, and the output visualized using the fineradstructureplot.r and finestructurelibrary.r R scripts (available at https://github.com/millanek/fineRADstructure). Populations were defined as clusters within the *fineRADstructure* tree and relatedness plot.

The results of the cluster analyses based on the ddRAD-seq dataset were compared with similar analyses suitable for the panel of STRs described above. Specifically, we performed cluster analyses on the STRs dataset using the Bayesian method implemented in STRUCTURE v2.3.4 (Pritchard, Stephens, & Donnelly, 2000). Analyses were conducted choosing a model with admixture, uncorrelated allele frequencies, and a non-uniform ancestry prior *alpha* among clusters, as suggested by (Wang, 2017) for uneven samplings. We ran 20 replicates for each value of K (i.e., the number of source populations) from K=1 to K=10 with 750,000 MCMC after a burnin of 500,000. Structure results were summarized and visualized with the web server CLUMPAK (Kopelman, Mayzel, Jakobsson, Rosenberg, & Mayrose, 2015). We used STRUCTURE HARVESTER (Earl & vonHoldt, 2012) to infer the best value of K, based on both deltaK (Evanno, Regnaut, & Goudet, 2005) and the Pr[X|K] (Pritchard et al., 2000).

We further tested for population structure both the ddRAD-seq and STR datasets using a Principal Component Analysis (PCA) with the *dudi.pca* function in the R package *adegenet* v2.1.3 (Jombart, 2008). The samples were visualized following the clusters inferred by *fineRADstructure*.

Pairwise population F_ST_ values were estimated among clusters inferred from *fineRADstructure* analysis using ARLEQUIN v3.5 (Excoffier & Lischer, 2010) for ddRAD-seq and STR datasets.

### 2.4 Design of a small panel of informative SNPs

We applied three different methods suitable to select informative SNPs useful for geographic assignment (see Helyar et al., 2011 for review): loadings from Principal Component Analysis (PCA), F_ST_, and random forest (RF) (Breiman, 2001). Our aim was to identify the best panel with 48 or 96 SNPs, meaning that a total of six panels were compared (two panel sizes for each method). Only SNPs having at least 50 bp flanking region in the ddRAD loci were selected, as required by the KASP screening protocol (see below). The ability of the panels to reflect the geographic assignment based on the complete set of SNPs was analyzed following a “node approach” based on the dendrogram produced by *fineRADstructure* with the whole dataset (see Results). This dendrogram represents the maximum level of geographic resolution supported by the entire set of SNPs. To maintain this level of resolution while reducing the number of markers, we considered one node at the time, and ranked each SNP with the aim of identifying SNPs that were most informative for maximizing genetic differentiation between the two groups separated by the node. A comparable number of SNPs was selected from each node (see Results).

*PCA-based panel*. We performed seven principal component analyses, one per node, using the R package *adegenet* v2.1.3 (Jombart, 2008). For each PCA, a graph was created to visualize the distribution of the data and evaluate which axes would better separate the distinct groups; then for each informative axis, we saved the SNPs within the top 5% loading value.

*F_ST_-based panel*. The R package *genepop* v1.0.5 (Rousset, 2008) was used to compute pairwise F_ST_ between the two groups branching from each node, retaining the SNPs with the highest F_ST_ values.

*Random forest panel*. We finally applied Random Forest (RF; R package *randomforest* v4.6-12; Liaw & Wiener, 2002), a machine learning method used for classification and regression. This method allows to identify a limited number of features (e.g., SNP) that best classify the observations into prior groups by computing the Mean Decrease in Accuracy (MDA). MDA is a measure of worsening of the model in classifying the observations when removing one feature at the time. Higher MDA means that the feature is important to the model. At every run, a forest of decision trees is grown. The out-of-bag (OOB) error is how RF measures misclassification; it is the mean of the prediction error of a bootstrapped training sub-sample across all the trees. Following the node approach as for the previous methods, we determined for every node our *ntree* parameter (number of trees) by running RF with 100, 500, 1,000, 8,000, 25,000 trees. The *mtry* parameter (number of features considered at each node) was tested using the function tuneRF. We then ran RF 10 times per node, as suggested in Sylvester et al. (2018), computed and exported MDA from every run. We only selected SNPs that were listed in all the runs per node with MDA > 1. We finally created two panels retaining SNPs with the highest MDA.

Venn diagrams (http://bioinformatics.psb.ugent.be/webtools/Venn/) were built to visualize overlapping SNPs selected across all methods within each of the two panel sizes.

We performed self-assignment tests to compare the three methods in terms of assignment accuracy. Assignment testing was performed by using the R package *AssignPop* (Chen et al., 2018). Initially, the data set is divided into training (baseline), and test (holdout) data sets using a resampling cross-validation approach by the function *assign.MC*. The user can specify the proportion of individuals from each source “population” to be used in the training data set. High grading bias is avoided using this method by producing randomly selected, independent training and test data sets (Anderson, 2010). Furthermore, using a PCA, the dimensionality of the training data sets (i.e., the genotypes) is decreased, and the result is utilized to create prediction models using user-chosen classification machine learning algorithms (Chen et al., 2018). Lastly, the models are used to estimate membership probabilities of tested individuals and assign them to a source population. Additionally, the training data are evaluated, and assignment tests are performed to assess the origin of individuals (Chen et al., 2018). Resampling was repeated 500 times for each combination of training individuals and loci. The proportion of individuals from each source population randomly allocated to the baseline data set was set to 0.5 and 0.7. Finally, the Linear Discriminant Analysis (LDA) was used as a classifier model.

### 2.5 Test and validation of the SNP panel

After evaluating the performances and the cost-effectiveness of the 96 and 48 SNP panels, we selected the smaller set of markers (48 SNPs) identified by the F_ST_ approach and these loci were tested with the KASP-by-Design Fluidigm Assays (LGC Genomics) (He et al., 2014). Using this panel, we typed a total of 190 DNA samples including samples previously typed with other markers (ddRAD-seq and/or 7 STRs;) as internal control, samples from captivity (from rescue centres), or from wild populations never typed before (see **Table S1** for a detailed description of the samples). Using the SNP data obtained with ddRAD-seq as a reference database (70 samples) we performed an assignment analysis with STRUCTURE (POPFLAG for individuals in the reference database; “update allele frequencies using only individuals with POPFLAG=1” option under a USEPOPINFO without admixture model). K was fixed at its optimal value (see results), while the run length and other parameters were set as above (see STRs analysis with STRUCTURE). We assigned individuals to a source population when the probability of an individual belonging to that population was above 80%. Exclusion tests were performed with the partially Bayesian exclusion method (Rannala & Mountain, 1997) implemented in GENECLASS2 (Piry et al., 2004). We compared observed genotypes with an expected likelihood distribution of genotypes generated for each reference population by simulating 1,000,000 individuals with Monte Carlo resampling. Furthermore, we performed assignment analysis using 7 STR loci on a subset of samples (68 samples) for which STR data were already available using the same methods and parameters used for the SNPs data. The STR database developed in Biello et al. (2021) was used as a reference. Populations not included in the ddRAD-seq experiment were removed (e.g., France).

**TABLE 1.** Mean percent assignment scores using AssignPop with 0.7 as proportion of training individuals and 500 repetitions of Monte-Carlo cross validation. 48 or 96 SNPs selected with PCA, F_ST_ or RF.

| | Method | Assignment score<br>(%Mean ± %SD) |
| --- | --- | --- |
| **48 SNPs** | PCA | 75.5 ± 7.1 |
| | $F_{ST}$ | 83.2 ± 7.9 |
| | RF | 73.3 ± 6.5 |
| **96 SNPs** | PCA | 94.0 ± 4.6 |
| | $F_{ST}$ | 88.5 ± 6.7 |
| | RF | 88.2 ± 6.2 |

## 3 RESULTS

### 3.1 Filtering and quality control

From the 84 individuals sequenced, 14 samples were excluded due to low coverage and high amount of missing data. For the remaining 70 individuals the sequencing coverage obtained per individual ranged from 89,897 to 412,840 with an average of 236,722 reads per sample (see **Table S2**). After filtering loci for missing data, by retaining markers that were genotyped in at least 75% of the individuals, we recovered 1,179 loci and 3,182 SNPs (*full dataset*).

**TABLE 2.** Cost per sample for 7 STR markers (Biello et al., 2021; Perez et al., 2014) and 41 SNPs (present work) for an increasing number of samples. STRs: fragment analysis quotation with or without PCR amplification (second and third column, respectively) by Macrogen Europe; KASP quotation by LGC Genomics, UK (October 2021). For KASP a minimum of 22 samples is required to determine the genotype of all the data points by means of genotyping cluster generation.

| Number of samples | 7 STRs (in-house PCR, outsourced fragment analysis) | 7 STRs (outsourced PCR and fragment analysis) | 41 SNP panel (outsourced KASP, LGC genomics) |
| --- | --- | --- | --- |
| 12 | 23.3€ | 34.4€ | - |
| 24 | 23.3€ | 34.4€ | 14.2€ |
| 48 | 23.3€ | 34.4€ | 10.6€ |
| 96 | 18.8€ | 30.1€ | 9.4€ |
| 192 | 18.8€ | 30.1€ | 7.0€ |

### 3.2 Genetic structure with ddRAD-seq and comparison with STRs

The co-ancestry matrix obtained with *fineRADstructure* using the *full dataset* showed the presence of eight main clusters (**Figure 1a**). The lowest amount of co-ancestry was found between the two subspecies, *T. h. boettgeri* and *T. h. hermanni*. Within the *T. h. hermanni* group, six distinct clusters were identified: Italian Peninsula (ITP), North Calabria (NCA), Central Calabria (CCA), South Calabria (SCA), Sicily (SIC) and Sardinia (SAR). These results support a finer genetic structure than previously suggested using STRs (Biello et al., 2021; Perez et al., 2014). More specifically, SNPs allowed the identification of three clusters within the Calabria region (NCA, CCA, SCA) and a distinction between Sicily and Sardinia (SIC, SAR) (**Figure 1b**). The average co-ancestry between the six clusters of *T. h. hermanni* was generally low, with the higher co-ancestry levels found between ITP and NCA, and between SIC and SAR. Within the *T. h. boettgeri* group, we detected two clusters, namely Greece (GRE), and Bosco Mesola + Croatia (MEC). The comparison between individuals within these two genetic clusters showed the highest level of co-ancestry compared to other groups.

**FIGURE 1.**
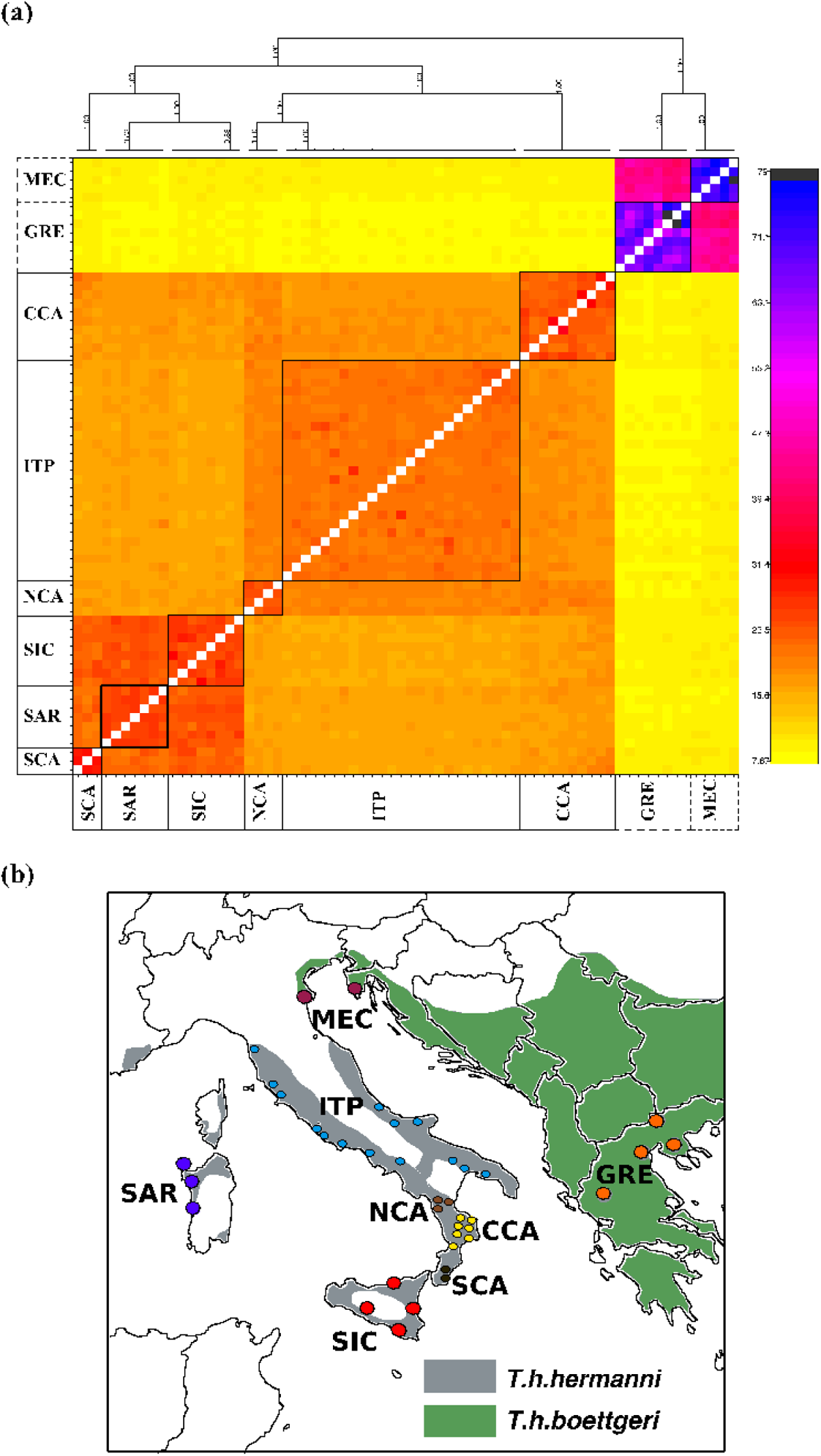
(a) Clustered *fineRADstructure* co-ancestry matrix. The analysis included individuals from 8 areas (GRE=Greece; MEC=Croatia and Bosco Mesola; ITP=Italian Peninsula; NCA=Northern Calabria; CCA=Central Calabria; SCA=Southern Calabria; SIC=Sicily; SAR=Sardinia) and two subspecies (*Testudo hermanni hermanni*: solid line, *Testudo hermanni boettgeri*: dashed line). The highest levels of co-ancestry are indicated in black/purple and the lowest in yellow. (b) Geographic distribution of the sampled populations.

Population clustering was also investigated by means of principal components analysis on the independent SNPs (the *single SNP dataset*). The first three principal components, PC1, PC2 and PC3, together accounted for 52% of the total variance in the dataset, with eigenvalues of 39.4%, 8.4% and 4.2% respectively (**Figure 2a,b**). PC1 separated the two subspecies, whereas PC2 separated the western subspecies *T. h. hermanni* in two groups, the first composed by three of the four mainland Italian populations (ITP, NCA, and CCA) and the second by south Calabria together with the two islands (SCA, SIC, and SAR) (**Figure 2a**). Within the first group, ITP, NCA and CCA were distributed along a “north-to-south” gradient, as were the corresponding regions of origin of the populations; conversely within the second group no clear spatial structure could be identified. The third axis separated the two *T. h. boettgeri* clusters, GRE and MEC (**Figure 2b**).

**FIGURE 2.**
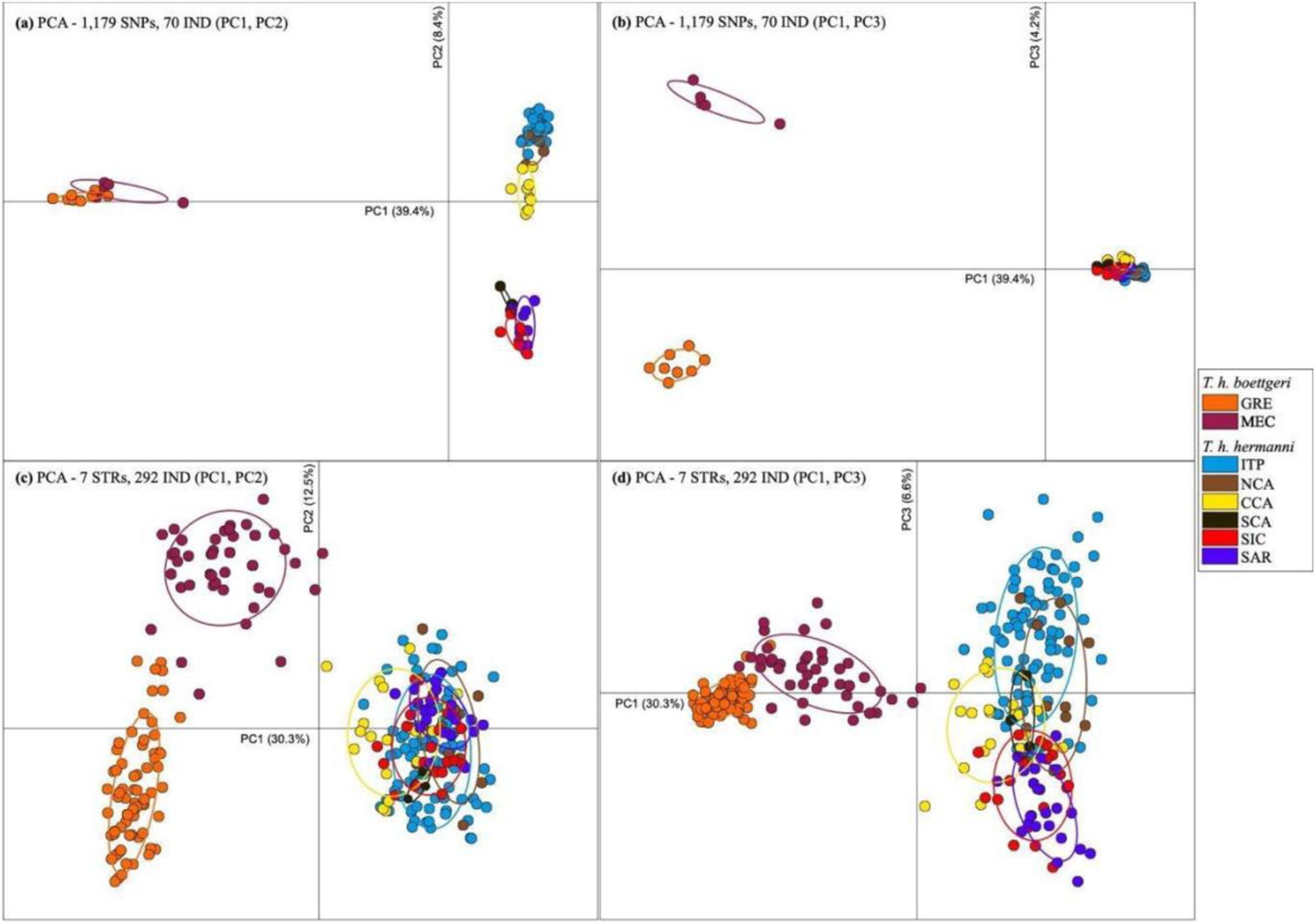
Principal component analysis (PCA) of (a, b) 1,179 SNPs (from ddRAD-seq) with 70 samples, and (c, d) 7 STRs with 292 samples. Coloured by eight groups found with *fineRADstructure* (see Figure 1). GRE=Greece; MEC=Croatia and Bosco Mesola; ITP=Italian Peninsula; NCA=Northern Calabria; CCA=Central Calabria; SCA=Southern Calabria; SIC=Sicily; SAR=Sardinia.

Pairwise estimates of F_ST_ among western subspecies populations ranged from 0.05 (between SIC and SAR) to 0.35 (between the ITP and SCA) (**Figure S1**, lower diagonal). Within the two subspecies, the highest F_ST_ value was observed between the two eastern subspecies populations MEC and GRE, (F_ST_ = 0.47). When populations belonging to different subspecies are compared, F_ST_ values reached values between 0.7 and 0.8. All F_ST_ values were significant at the nominal significance level of 0,05, and most P-values were lower than 0.01 (see **Table S3**).

The SNPs markers isolated with the ddRAD-seq approach appeared thus able to genetically discriminate populations with different geographic distribution, with a precision that was not the same using different statistical approaches. To further confirm that the SNPs markers contained relevant information to discriminate individuals with different geographic origin, at least at the macro-areas level, we compared the geographic structure inferred with the SNPs with that estimated from a small STR panel of proven efficacy for conservation and forensic applications (Biello et al., 2021). The goal here was not to analyze in detail the difference between the different markers, but to additionally validate the SNPs dataset (based on 70 individual and >1000 independent biallelic loci) by showing that it contains the same geographic partition of the genetic variation inferred from a different type of molecular marker (almost 300 individuals typed at 7 STRs). We can confidently conclude that the geographic structure identified by the SNPs is very similar, and slightly more resolved, than that based on a much large sample size because: i) the pairwise F_ST_ values between the groups inferred by the SNPs and the STRs (**Figure S1,** upper diagonal) were highly correlated (see **Figure S2**); ii) the PCA results on the STR dataset confirmed a clear subdivision of the two subspecies, as well as the two *T. h. boettgeri* clusters GRE and MEC (**Figure 2c,d**); overall, the first three components separated some of the *T. h. hermanni* populations, but with higher overlap between individuals, as observed in the SNPs dataset (**Figure 2a,b**); iii) the major genetic clusters identified by the STRUCTURE analysis on the STR markers (see **Figure 3** and **Figure S3**) corresponded to those observed with the SNPs dataset, but without separating the three pairs of areas with the smallest F_ST_ values (ITP and NCA; CCA and SCA; SIC and SAR).

**FIGURE 3.**
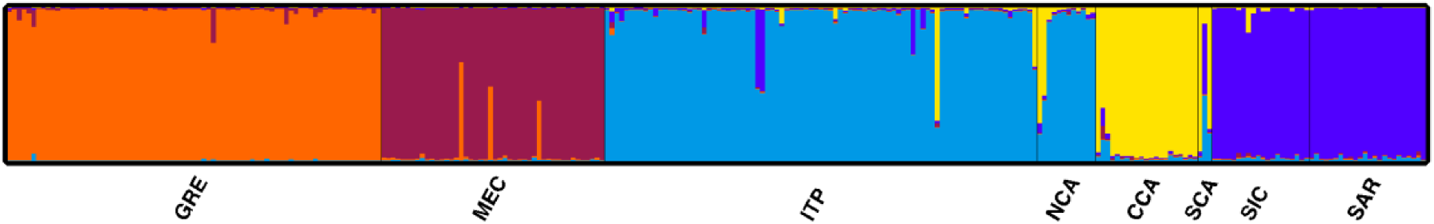
Genetic structure at seven STR loci of wild *Testudo hermanni* populations estimated using STRUCTURE using 292 samples and K=5 with 8 groups. GRE=Greece; MEC=Mesola and Croatia; ITP=Italian Peninsula; NCA=Northern Calabria; CCA=Central Calabria; SCA=Southern Calabria; SIC=Sicily; SAR=Sardinia.

### 3.3 Isolation of a small panel of informative SNPs

The SNPs were selected following a “node approach” based on the dendrogram produced by *fineRADstructure* with the whole dataset (**Figure 4**), with the panels including a comparable number of SNPs from each node.

**FIGURE 4.**
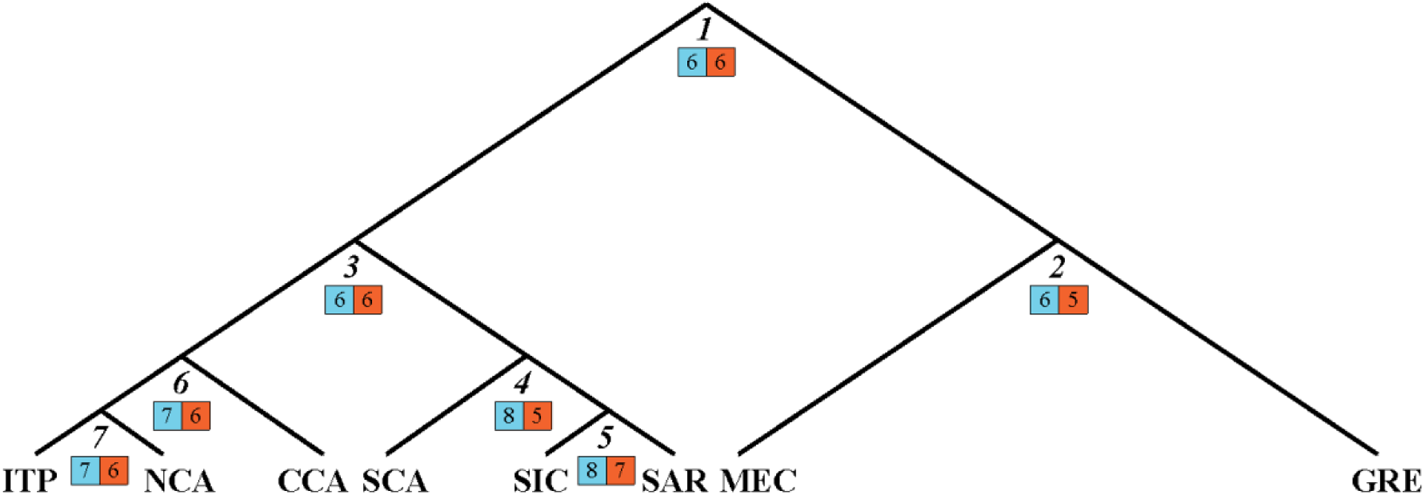
Genetic relationships among the clusters identified with *fineRADstructure* based on co-ancestry. The numbers in the squares indicate for each node the SNP included in the panel (left, light blue) and those successfully typed (right, orange). GRE = Greece; MEC = Croatia and Bosco Mesola; ITP = Italian Peninsula; NCA = Northern Calabria; CCA = Central Calabria; SCA = Southern Calabria; SIC = Sicily; SAR = Sardinia.

In the first method (PCA-based panel) the highest loadings from the top 5% of the most informative principal components (PC) were selected. For all the seven nodes, PC1 was always the only axis to highlight differences in our data. In the second method (F_ST_-based panel) the loci with the highest F_ST_ resulting from pairwise comparisons between the two clades at each node were selected. F_ST_ values ranged from 1 to 0.39 in the 48 SNPs panel and from 1 to 0.33 for the 96 SNPs panel. Finally, in the last method (RF) the loci with the highest MDA were included. MDA values ranged from 7.99 to 1.45 in the 48 SNPs panel and from 7.99 to 1.26 in the 96 SNPs panel. The distribution of SNPs across the three SNP selection methods is summarized by means of a Venn diagram in **Figure S4**.

The six different SNP panels (48 or 96 SNPs, three selection criteria) were tested for accuracy in providing the assignment of individuals to their actual group. Assignment scores (**Table 1**) were high and ranged from 73.3% to 94%.

When the 48 SNP panels were tested, individuals were assigned to their original cluster at a mean accuracy of 75.5%, 83.2% and 73.3%, for PCA, F_ST_ and RF methods, respectively (**Table 1, Figure S5**). Assigning individuals to the two subspecies was extremely accurate, ranging from 99.1% to 100% depending on the method used (see **Figure S6**). Accuracy was very high also when we attempted to assign individuals to the two main groups in *T. h. hermanni*, Italian peninsula (including ITP, NCA and CCA clusters) and the Mediterranean Islands (including SCA, SIC and SAR). The mean accuracy ranged from 91.6% (PCA) to 97.6% (RF) (see **Figure S7**).

The 96 SNPs panels assignments showed a mean accuracy of 94%, 88.5% and 88.2%, PCA, F_ST_ and RF methods, respectively (**Table 1, Figure S8**). Assigning individuals to the 2 subspecies was 100% accurate independently from the method used (see **Figure S9**). The mean accuracy when we assigned individuals to the two main groups in *T. h. hermanni* was extremely accurate, ranging from 98.5% (F_ST_) to 99.1% (RF) (see **Figure S10**).

### 3.4 Test and validation of the SNP panel

Considering the relatively low power increase obtained by doubling the size of the SNPs panel, but the relevant additional costs of the larger compared to the smaller panel (see Discussion), we decided to empirically test the smaller panel that has a greater chance to be incorporated in future large scale genetic testing in this species.

Of the 48 SNPs initially selected with the F_ST_ approach, seven were dropped due to technical problems. Thus, 41 SNPs were left for further assignment analysis.

The genotypes of 189 DNA samples (one sample failed) were assigned to the most probable geographic area of provenance using STRUCTURE and the ddRAD-seq database of 70 samples as reference. We used K=8 and membership coefficients (Q-value) > 0.80 as the assignment threshold. Using these parameter values, we were able to assign more than 78% of samples to one of the eight clusters. 32.8% of the tortoises were assigned to the GRE cluster, 24.3% to ITP, 18% to MEC, 2.1% to the SAR, 0.5% to NCA and 0.5% to CCA, while 21.7% of the individuals were not assigned to any predefined cluster (NA).

Furthermore, we compared the geographical assignment of samples that were genotyped at 41 SNPs and 7 STR loci, using the SNP reference (see **Figure 4**) and STR reference (see Biello et al., 2021), respectively. In order to minimize the differences between the two references we decided, in this analysis, to group together the clusters SIC and SAR inferred by SNPs following the STRs cluster ISS (see Biello et al., 2021). Assigning individuals to their area of origin was extremely accurate when we considered samples from wild populations. Both markers assigned all the samples to the same cluster (**Figure 5**). In contrast, for the individuals from captive populations, the results were not completely concordant between the two markers. Approximately 83% of samples were assigned to the same groups, while 17% of samples were assigned to different clusters or to a cluster only with one of the two panels (**Figure 5**). Among the unassigned samples, four samples assigned to GRE with the SNPs were assigned to MEC with the STR data (clusters from the same subspecies, *T. h. boettgeri*), and three samples assigned to ITP with the STRs were assigned to ITP and CAL with the SNPs (clusters from the same subspecies and adjacent geographic regions, *T. h. hermanni*).

**FIGURE 5.**
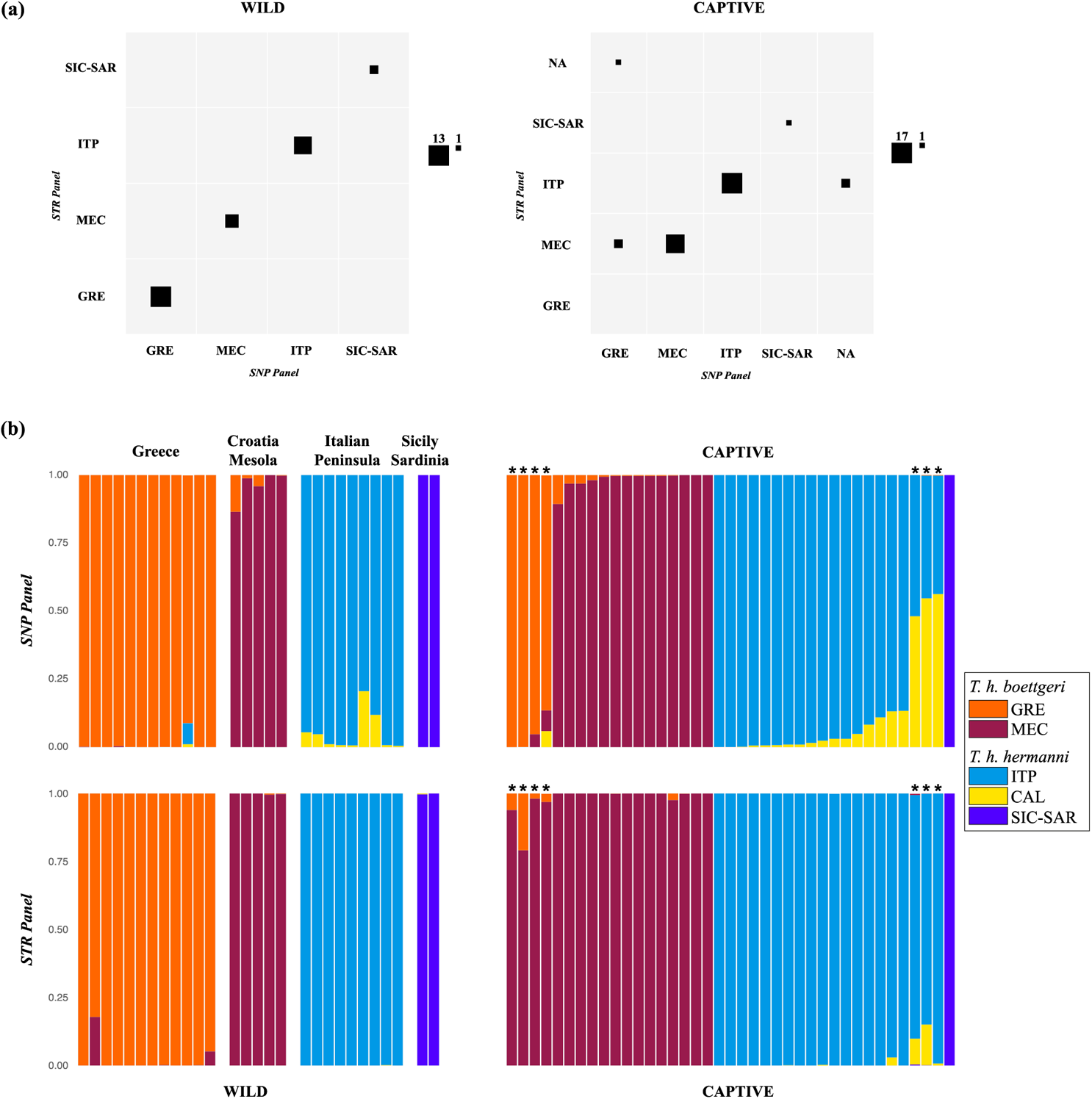
Contingency tables (a) and bar plots (b) of the geographical assignment of 67 samples, 28 Wild (left) and 39 Captive (right), characterized at 41 SNPs and 7 STR loci, to the SNP reference (see Figure 5) and STR reference, respectively. a) Columns correspond to the number of individuals assigned to original clusters using the SNP panel, while rows correspond to the number of assignments using the STR panel (Biello et al., 2021): square sizes are scaled by number of assignments to the cluster. SNP/STR Panels: GRE = Greece; MEC = Croatia and Bosco Mesola; ITP = Italian Peninsula; SIC-SAR = Sicily and Sardinia; NA = Not Assigned. * Individuals assigned to different clusters by the two panels.

Assigning individuals from Italian rescue centres that were genotyped for the first time showed evidence of long-distance translocations and potential hybridization events (**Figure 6**). For example, 46% of the tortoises from the Veneto center were not assigned to any cluster, 27% were classified as imported from Greece (GRE), 19% were classified within the *T. h. hermanni* subspecies (Italian Peninsula, ITP), and only 8% were assigned to the MEC cluster (typically found in North-Eastern Italy). In the Emilia-Romagna center, 58% of the tortoises had Greek origins, and 6% were assigned to the MEC cluster. Approximately one-third of the tortoises from this centre was not assigned. The centres of Emilia-Romagna and Lazio show a similar proportion of unassigned individuals (33% and 22%, respectively). On the contrary, the two centres show an opposite trend in terms of individuals assigned to the GRE (i.e., non-native origin) and ITP (i.e., local origin) clusters: 58% GRE and 6% ITP for the Emilia-Romagna centre, 11% GRE and 56% ITP for the Lazio center.

**FIGURE 6.**
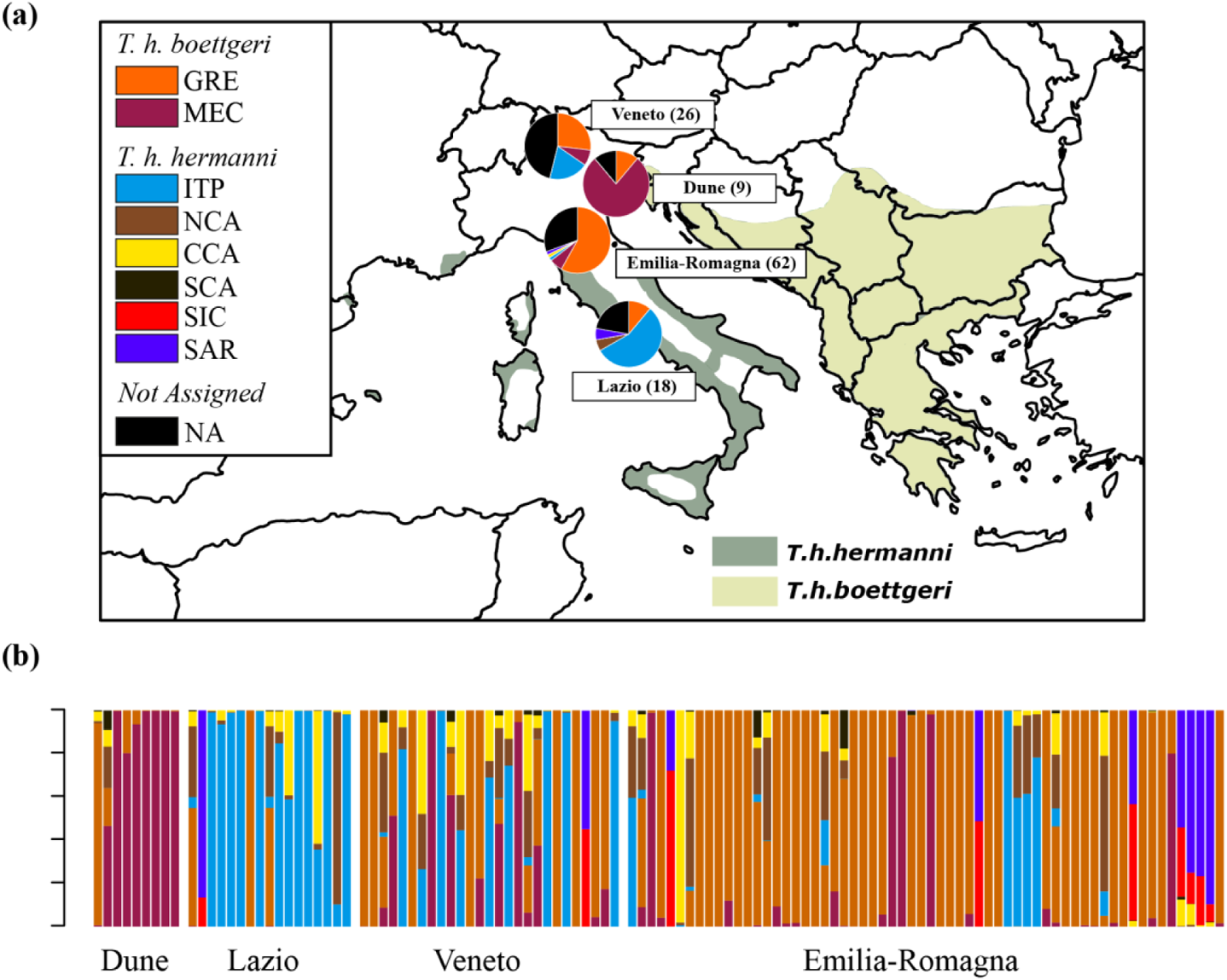
Geographic assignment of 115 confiscated samples from three Italian rescue centres (Veneto, Emilia-Romagna and Lazio) and one wild population (Dune). Local assignments for each location are showed in the pie charts on the map (in brackets the samples size). GRE = Greece; MEC = Croatia and Bosco Mesola; ITP = Italian Peninsula; NCA = Northern Calabria; CCA = Central Calabria; SCA = Southern Calabria; SIC = Sicily; SAR = Sardinia; NA = not assigned samples.

Interestingly, the tortoises from a wild and isolated population located in a natural reserve with fossil dunes corresponding to the former coastline of the Northern Adriatic Sea (Massenzatica Reserve), sampled here for the first time, were mainly assigned to the MEC cluster (78%). Populations from this cluster are typical in nearby areas (North-East Italy), showing a likely native component of this population.

## 4 DISCUSSION

In this study, we present the development and the application of a small panel of SNP markers useful to investigate population genetic structure and geographical assignment in the endangered species Hermann’s tortoise, *Testudo hermanni*. We initially performed a ddRAD-seq experiment on 70 individuals from different locations, and we showed that these loci, in terms of genetic structure and power to identify the macro-area of origin, outperform those obtained with a sample of 292 wild individuals previously typed at 7 STRs. Starting from these results, we designed a small panel of SNPs, retaining the most informative markers in terms of maximizing F_ST_ between the clusters inferred by the population genetic analysis. Finally, we tested the small SNP panel in 190 individuals using the KASP genotyping chemistry, a relatively inexpensive SNP genotyping approach. This cost-effective molecular tool enables a large number of confiscated individuals to be genotyped for geographical assignment, necessary for their management and their possible re-allocation in the wild.

### 4.1 Population genetic structure of *T. hermanni*: from STRs to SNPs

The overall results of the genetic structure of *T. hermanni* wild populations using SNPs confirmed the pattern based on the analysis of 7 STR loci found by Biello et al. (2021) but also revealed a more detailed structure. First, SNPs data were able to identify three genetically distinct groups in Calabria, a small region in the South of Italy, corresponding to samples collected in Northern, Central, and Southern areas, respectively. This result is not surprising given that this geographically heterogenous region is a hotspot of genetic diversity for many temperate species and likely acted as a single or multiple glacial refugia (Canestrelli, Aloise, Cecchetti, & Nascetti, 2010; Chiocchio, Bisconti, Zampiglia, Nascetti, & Canestrelli, 2017). Considering however that the samples are not covering homogenously the whole region, and the clustering algorithm we applied tend to overestimate genetic structure under isolation by distance (Frantz et al., 2009), we cannot exclude that these clades do not refer to genetic isolates but to genetically distinct groups sampled within an area of large genetic variation but limited barriers and genetic discontinuities. Second, SNP data suggest, differently from STRs, the existence of different genetic pools in Sicily and Sardinia. Finally, we note that even with a descriptive analysis such as the PCA, the SNPs data set produces groups more easily distinguishable from each other, supporting the higher discriminatory power for these markers when compared to seven STRs. In summary, even if the samples present in the ddRAD-seq dataset included only 23% of the individuals included in the STR dataset, these were sufficient to understand better the genetic structure in this species, especially in emphasizing the fine-scale structure among wild populations.

### 4.2 Design and validation of the SNP panel

Custom species-specific SNP panels, including high-ranking loci, have been shown to be highly informative for assignment studies (Förster et al., 2018; Jenkins, Ellis, Triantafyllidis, & Stevens, 2019; Kleinman-Ruiz et al., 2017) and prove to be an important resource for identifying the geographic origin of species in a conservation and forensic perspective.

We tested different methods for selecting highly informative SNPs and assuming that the target of the SNP number in the panel should be low to reduce costs. Using a panel of 48 SNPs, the F_ST_ method outperformed the Random Forest and PCA. The F_ST_ panel had a consistent higher self-assignment accuracy compared to those selected by PCA and RF. However, self-assignment of the largest panel (96 SNPs) showed a better accuracy of PCA method over F_ST_ and RF. This pattern is in discordance with what was found in Sylvester et al. (2018), where RF-based panels almost always outperformed F_ST_-based panels. The higher accuracy of the F_ST_ panel over the RF panel may be due to the fact that genetic structure in *T. hermanni* species is higher than in salmon populations, resulting in a higher fixation index (Sylvester et al., 2018). Interestingly, approximately half of the loci identified by each method in the 48 SNP panel are not shared among methods (approximately one third in the 96 SNPs panel), indicating that algorithms optimized the selection in very different ways (although the results were not dramatically different).

Given the relatively high overall self-assignment accuracy of the 48 SNPs panel based on F_ST_ (above 80%) and considering that the genotyping cost for a panel with twice as much of SNPs implied a two-fold cost increase, we selected the smallest panel for further genotyping to achieve a trade-off between accuracy and costs.

For the genotyping of our SNP panel, we adopted the SNP typing platform, KASP-by-Design Fluidigm Assays (LGC Genomics) (He et al., 2014). The SNP panel we have developed is extremely cost-effective: users only need to extract less than 500 nanograms of DNA (10 ng per sample per SNP for a genome size ranging from 2.0 to 3.5 Gbp) from any tissue and send it to the genotyping service, which will return easy-to-interpret table format output files. The genotyping cost is affordable, and data for 190 samples is provided for 1,338 € (7 € per sample). In comparison, for STRs, genotyping costs include fluorescently labeled oligos, PCR chemistry and hot start polymerase, with subsequent fragment analysis on a capillary sequencer. Since there are currently no multiplex PCRs for the 7 *T. hermanni*’s STRs, outsourcing the entire service is extremely expensive (**Table 2**). One option to reduce the cost of STR analysis is to carry out part of the wet lab internally, although KASP remains much cheaper (**Table 2**).

### 4.3 Test of the SNP panel

We observed a high genotyping success rate when we tested the panel with 190 samples, even with a low DNA yield (500 ng), which is significant in the case of noninvasively collected samples. We showed that missing data was below 1.9% for every SNP in our final panel, which is low compared with higher rates of missing data observed especially in STR genotyping of species of conservation concern where the sampling should not be invasive (Kraus et al., 2015). A few SNPs (seven) failed due to technical problems, most probably caused by non-specific annealing of primers in multiple genomic locations. The 41 SNPs performed very well in terms of individual identification. Assigning individuals to their area of origin was highly accurate (100%) when we considered samples from wild populations. These results were also confirmed by using a panel of STR markers (Biello et al., 2021). This is in accordance with previous studies, which showed that reduced panels of highly informative SNPs (<100 SNPs) performed as well as or better than the traditional number (10–20) of STR loci used for individual identification (e.g., Glover et al., 2010). We also showed that assigning individuals from captive populations to their location of origin was not completely concordant between the two markers. Approximately 17% of samples were not assigned to the same groups or not assigned to any group. This samples could represent descendent of crosses in captivity between individuals with different origin (including hybrids between sub-species), or to the lack of major genetic clusters in the reference populations.

Among the tortoises with unknown origin that we genotyped for the first time and are currently hosted in Italian rescue centres, we found evidence of long-distance translocations of individuals from Balkan regions and individuals with unclear genotype (i.e., potentially hybrids). This pattern confirms the Balkan areas as a source of illegal trading and the presence of hybrids in the Italian rescue centres due to mating occurring in captivity between individuals with different origins (see Biello et al., 2021).

Most of the tortoises from the wild population Riserva Naturale Orientata Dune Fossili Di Massenzatica (Emilia-Romagna, Italy), which was never sampled before, were assigned to the genetic cluster presents also in other areas in North-East Italy (Bosco Mesola) and Croatia. However, we also found non-native tortoises belonging to genetic cluster observed in Greek individuals.

### 4.4 Implications for management and perspectives

Approximately 2 million tortoises were exported from the former Yugoslavia to different European countries in the last century, especially after the second World War and until the 80’s (Ljubisavljević, Džukić, & Kalezić, 2011). A large fraction of them were *Testudo hermanni* harvested for the pet trade, and Italy was a major destination and virtually the only one where local populations already existed and introgression of non-native genomes could have occurred. Beside the possible translocation in recent and historical times of native individuals across the Italian peninsula and the Mediterranean islands (Perez et al., 2014), several thousands of individuals of Eastern origin, and their descendants, are therefore likely present in Italy in captivity (private and public seizures, including those with confiscated animals) and, probably at low frequencies (Biello et al., 2021; Perez et al., 2014), in some wild population. Considering that this species is endangered with a very patchy natural distribution in Italy (as well as in Spain and France), the possibility to monitor the genetic composition of wild populations and to select genetically suitable individuals for reintroduction projects should be carefully evaluated as a necessary step in conservation plans.

In a previous paper we introduced a panel of 7 STR markers useful to reach this goal (Biello et al., 2021). It was an important advancement for the assignment of individuals of unknown origin, based on a large reference dataset of 461 wild individuals collected across most of the distribution range of this species. Here we showed a new development based on a preliminary ddRAD-seq genomics study that allowed the introduction of an informative and cost-effective panel of biallelic markers suitable for the *Testudo hermanni* protection. This study, describing in detail the necessary methodological steps of the process, represents also a useful example for the development of similar tools in other species with similar management needs.

The SNP panel introduced in this study allows the genotyping of a large number of samples at low-cost. Compared to the development and the typing of a set of similarly informative STR markers, this approach is cheaper, faster, requires less handling, and provides immediate standardization across laboratories. At the moment, however, the reference database of wild populations for the SNPs panel is still smaller and geographically less representative of the distribution range of this species compared to the STRs database, and we therefore recommend, until more wild individuals will be typed, the use of the SNPs panel integrated for difficult assignments with the analysis of the STR markers.

Hundreds of tortoises are currently maintained in captivity in breeding and rescue centers, providing a highly valuable source of individuals for reintroduction and reallocation projects. A fraction of these individuals, selected considering their health conditions, sex, age, and genetic compositions, could represent not only a demographic and genetic supplementation of small natural populations, but also the founders of new wild populations in ecologically suitable areas where this species was present in the past. These interventions would favor the well-being of the captive tortoises but also the human well-being by creating more natural and biodiverse environments (e.g., Kuo, 2015; Fuller et al., 2007).

## ACKNOWLEDGMENTS

We thank Parco Natura Viva (Verona, Italy), Maurizio Bellavista, Michele Capasso, Giovanni Nobili, Fabrizio Bellini and all the Carabinieri Forestali at Punta Marina (Ravenna) for helping and providing samples from captive populations. We also thank Emanuele Lubian, Stefano Mazzotti and Annalaura Mancia for their help in collecting the specimens from Riserva Naturale Orientata Dune Fossili Di Massenzatica (Ferrara, Italy). We also thank Noemi Giurintano, Camilla Broggini and Maria Luisa Boglino for their help in laboratory work. This study was funded by the University of Ferrara, the Tuscia University (Viterbo) and the Carabinieri Forestali.

## CONFLICT OF INTEREST

None declared.

## DATA AVAILABILITY STATEMENT

Raw demultiplexed sequences are available on the Sequence Read Archive (SRA) on the study accession number: PRJNA777783.

## Notes

### Competing Interest Statement

The authors have declared no competing interest.

