## Supplementary Figure for "From STRs to SNPs via ddRAD-seq: geographic assignment of confiscated tortoises at reduced costs"

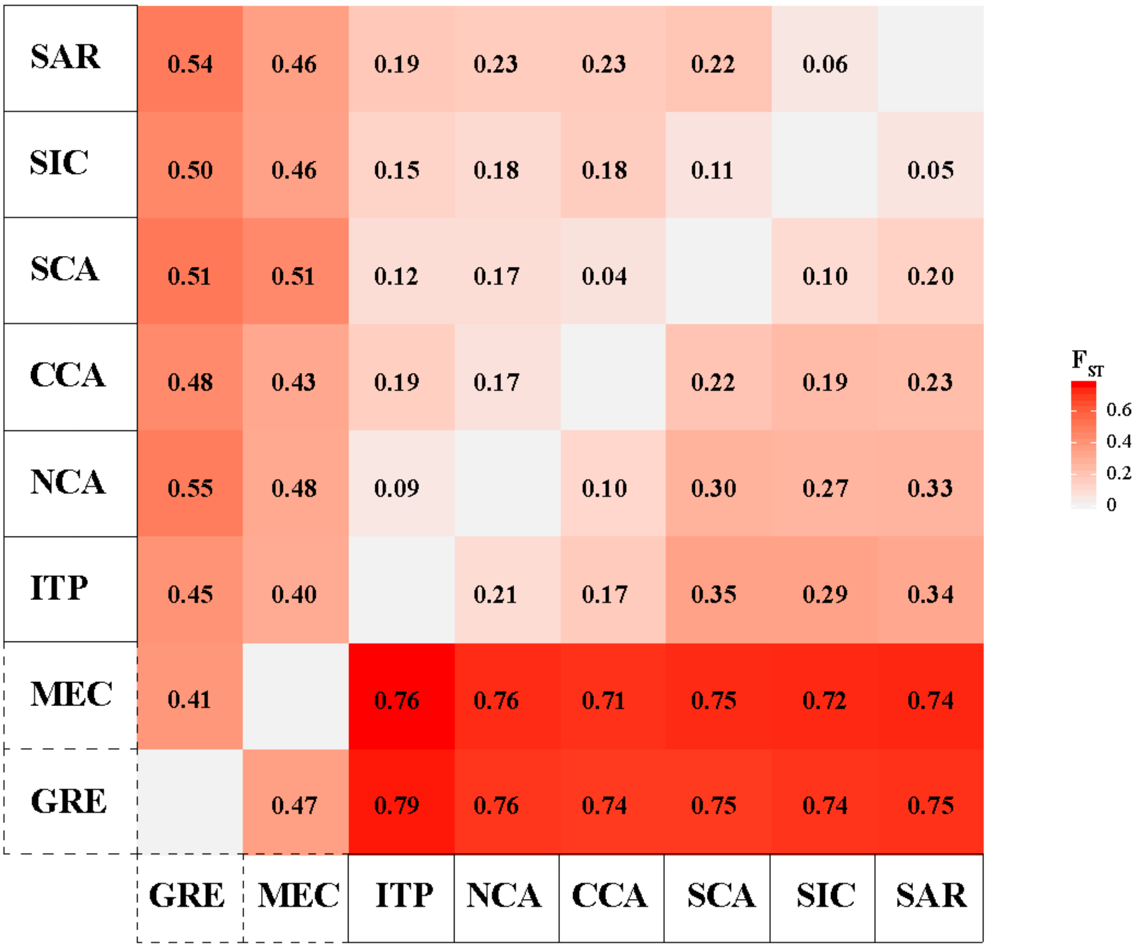


**FIGURE S1**. $F_{ST}$ values. SNPs (from ddRAD-seq) (below diagonal) and STRs (above diagonal). GRE=Greece; MEC=Mesola and Croatia; ITP=Italian Peninsula; NCA=Northern Calabria; CCA=Central Calabria; SCA=Southern Calabria; SIC=Sicily; SAR=Sardinia. The two subspecies *Testudo hermanni hermanni*: solid line, *Testudo hermanni boettgeri*: dashed line.

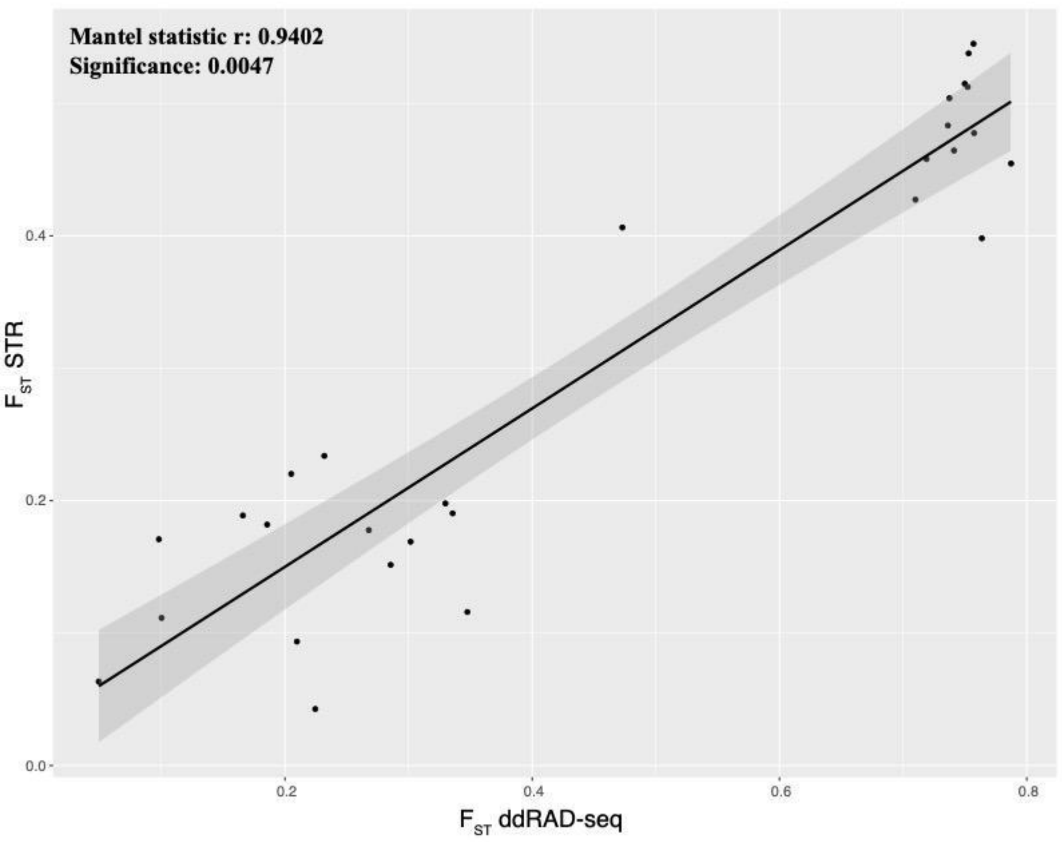


**FIGURE S2**. Mantel test comparing $F_{ST}$ values between SNPs (ddRAD-seq) and STRs datasets.

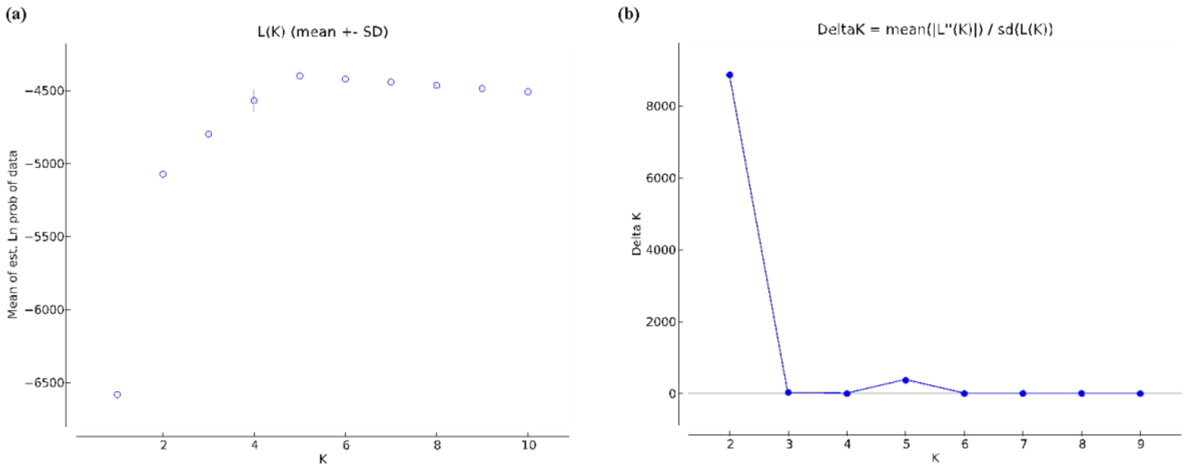


**FIGURE S3**. Probability of number of clusters (K) for 292 samples of *Testudo hermanni*, analysed at 7 STRs loci, using STRUCTURE HARVESTER (Earl et al., 2012). The Ln likelihood value (a) as described by Pritchard et al (2000) and the Delta K method (b) by Evanno et al. (2005).

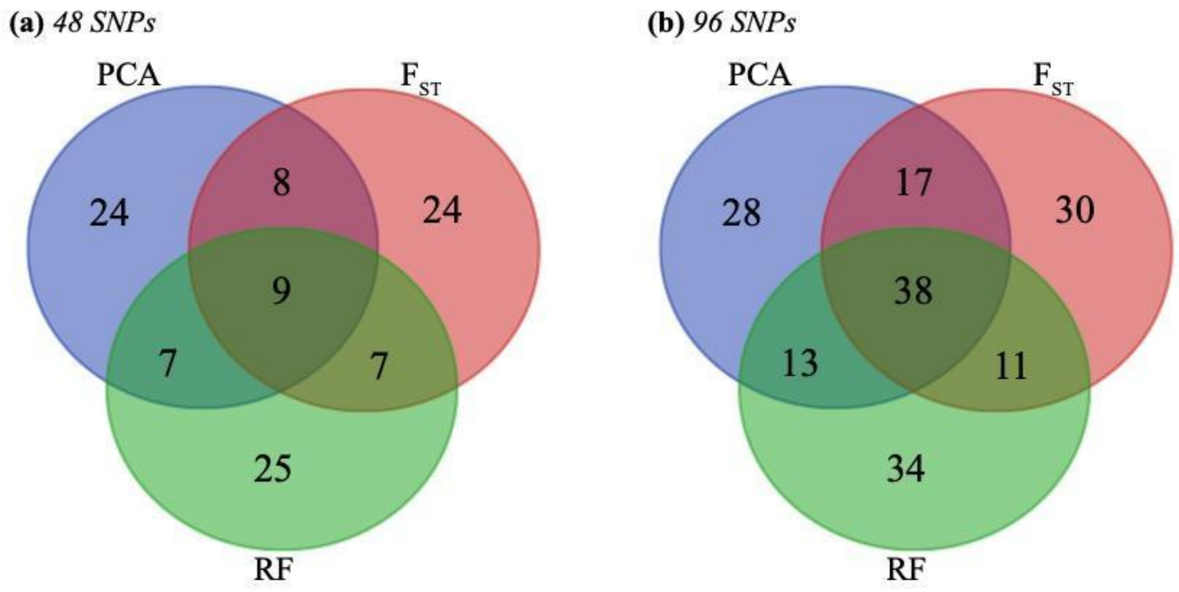


**FIGURE S4**. Venn diagram of SNPs shared among panel classes of 48 SNPs (a) and 96 SNPs (b).

**(a)** PCA

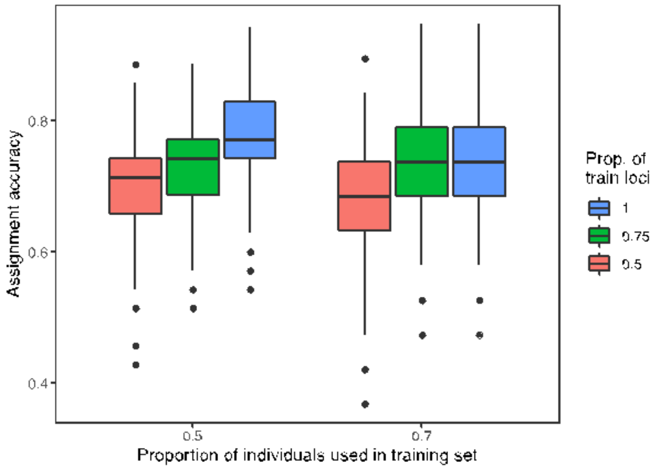


**(b)** $F_{ST}$

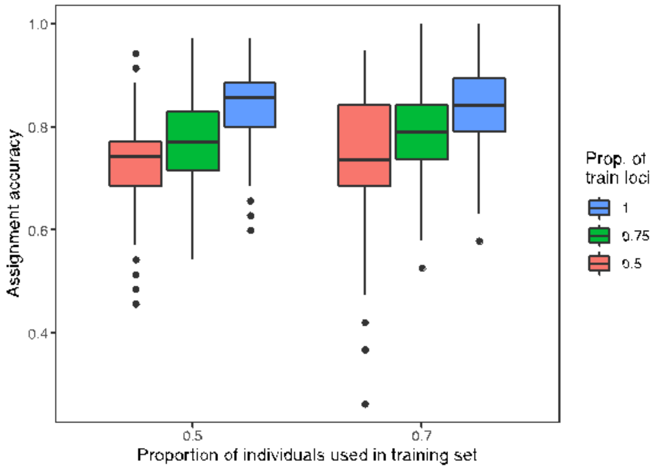


**(c)** RF

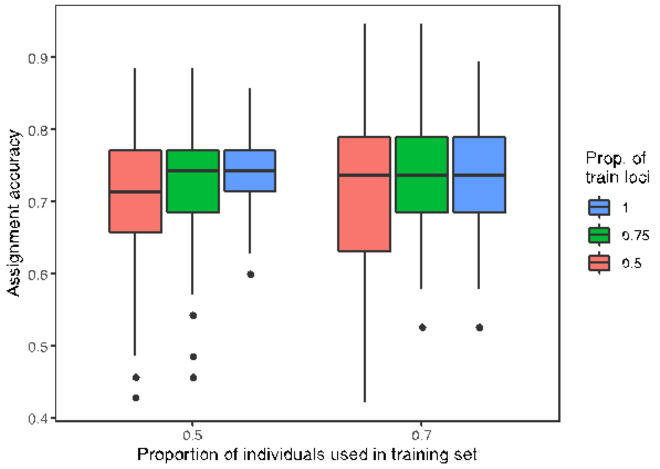


**FIGURE S5.** Overall assignment accuracies to the eight genetic clusters estimated via Monte Carlo cross-validation of the 48 SNPs panels, with two levels of training (baseline) individuals (50% and 70% of individuals from each group) crossed by up to three levels of training loci (top 50%, 75% and all loci) by 500 resampling events: (a) PCA; (b) $F_{ST}$; and (c) RF.

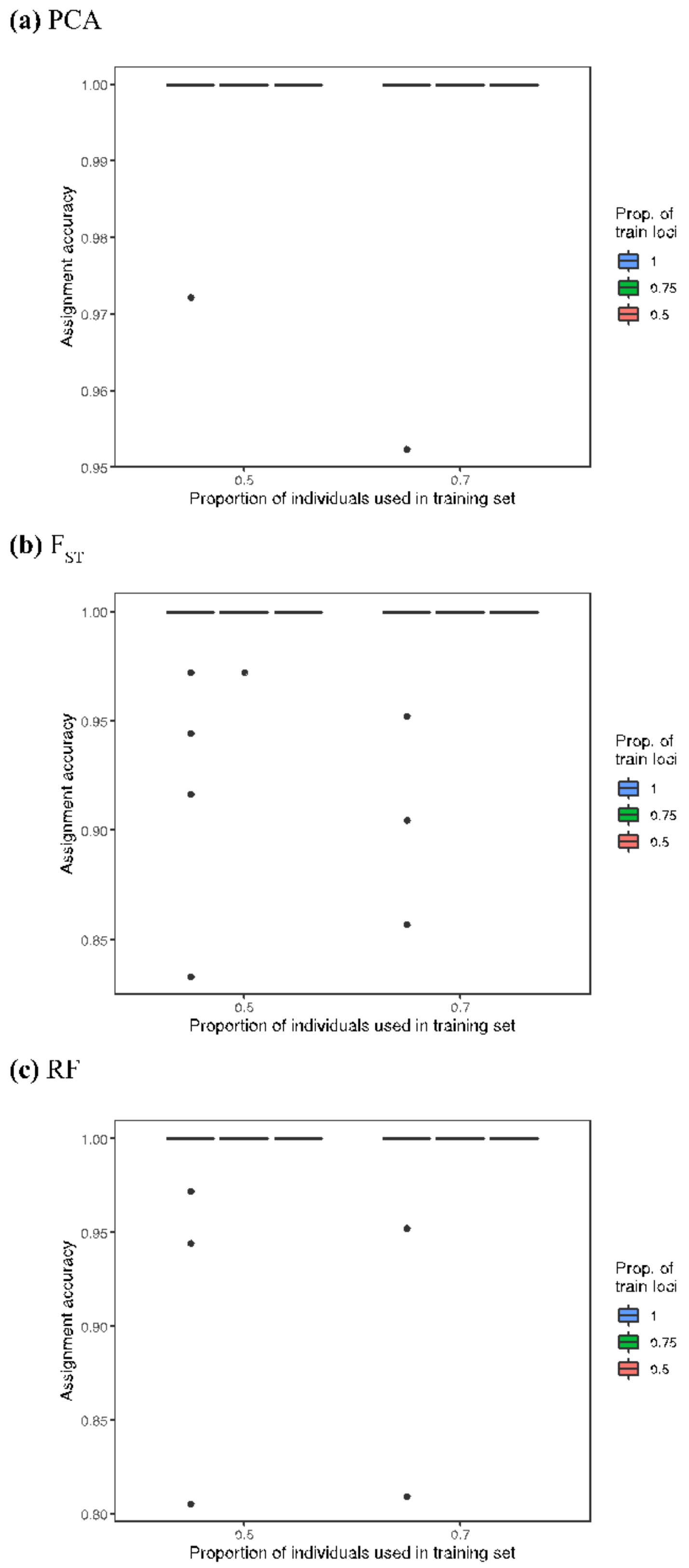


**FIGURE S6**. Overall assignment accuracies to the two subspecies (*T. h. boettgeri* and *T. h. hermanni*) estimated via Monte Carlo cross-validation of the 48 SNPs panels, with two levels of training (baseline) individuals (50% and 70% of individuals from each group) crossed by up to three levels of training loci (top 50%, 75% and all loci) by 500 resampling events: (a) PCA; (b) $F_{ST}$; and (c) RF.

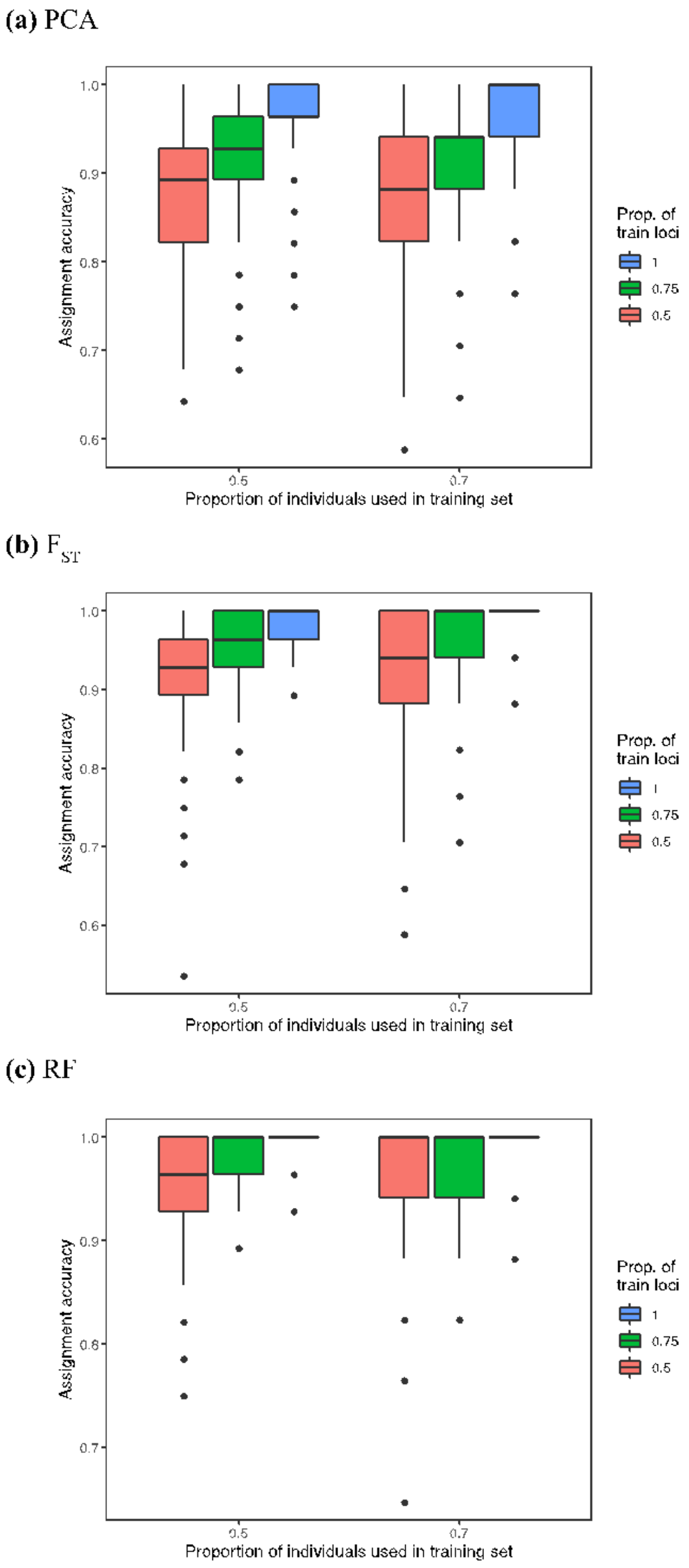


**FIGURE S7**. Overall assignment accuracies to the two main *T. h. hermanni* groups, Italian Peninsula (ITP, NCA and CCA) and Mediterranean Islands (SCA, SIC and SAR), estimated via Monte Carlo cross-validation of the 48 SNPs panels, with two levels of training (baseline) individuals (50% and 70% of individuals from each group) crossed by up to three levels of training loci (top 50%, 75% and all loci) by 500 resampling events: (a) PCA; (b) $F_{ST}$; and (c) RF.

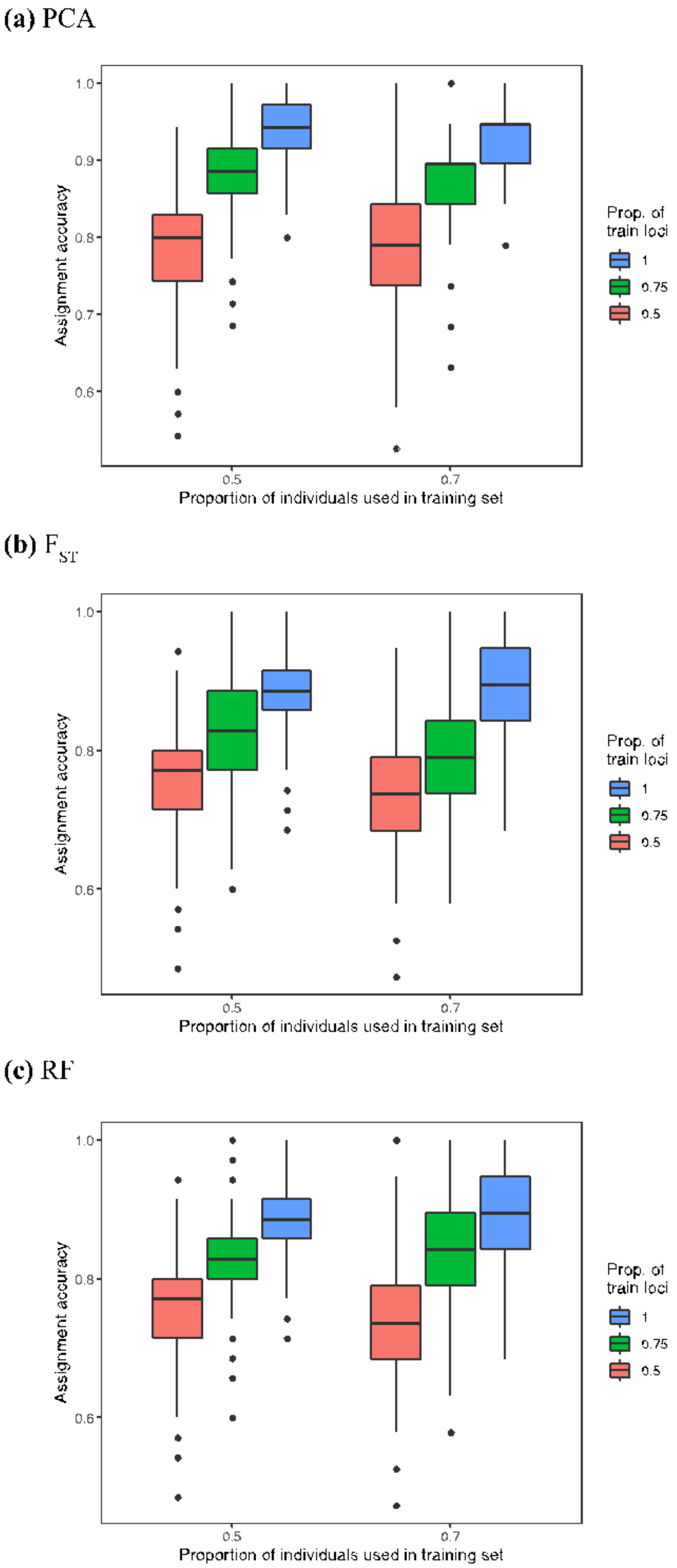


**FIGURE S8**. Overall assignment accuracies to the eight genetic clusters estimated via Monte Carlo cross-validation of the 96 SNPs panels, with two levels of training (baseline) individuals (50% and 70% of individuals from each group) crossed by up to three levels of training loci (top 50%, 75% and all loci) by 500 resampling events: (a) PCA; (b) $F_{ST}$; and (c) RF.

**(a)** PCA

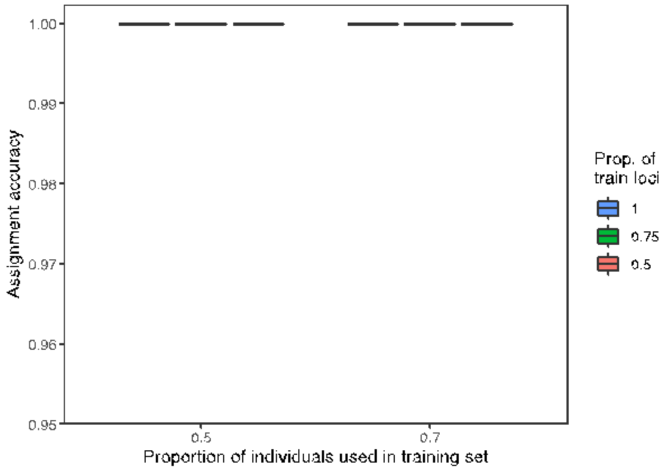


**(b)** $F_{ST}$

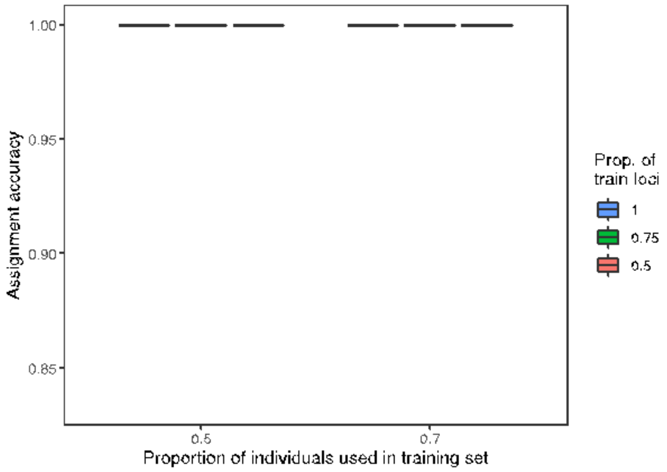


**(c)** RF

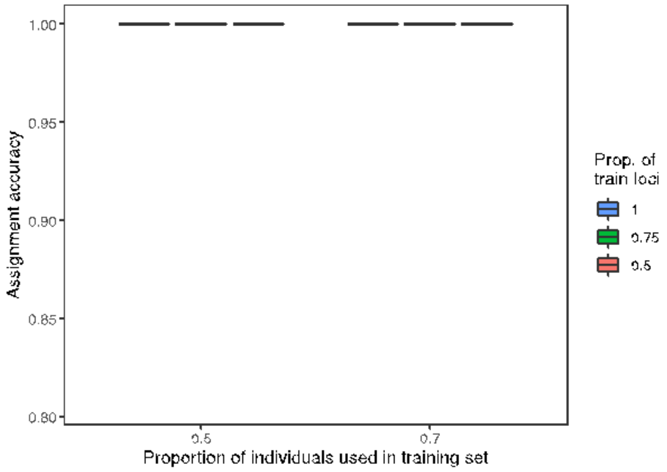


**FIGURE S9**. Overall assignment accuracies to the two subspecies (*T. h. boettgeri* and *T. h. hermanni*) estimated via Monte Carlo cross-validation of the 96 SNPs panels, with two levels of training (baseline) individuals (50% and 70% of individuals from each group) crossed by up to three levels of training loci (top 50%, 75% and all loci) by 500 resampling events: (a) PCA; (b) $F_{ST}$; and (c) RF.

**(a)** PCA

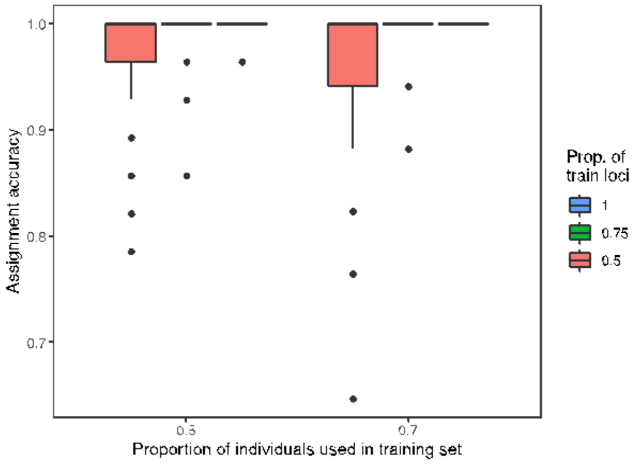


**(b)** $F_{ST}$

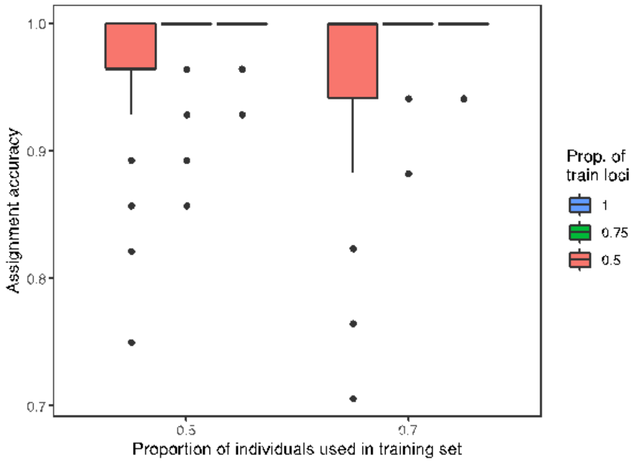


**(c)** RF

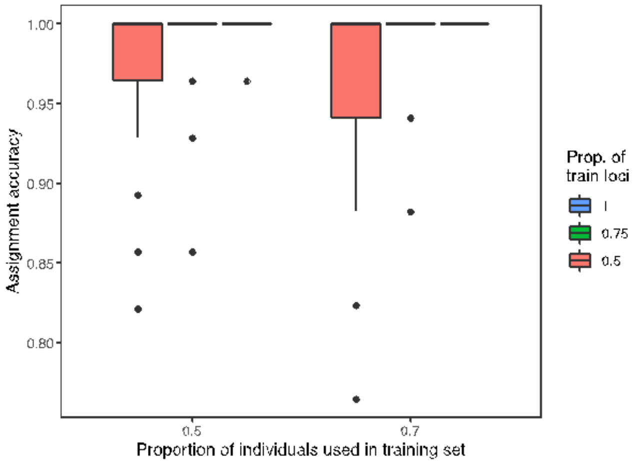


**FIGURE S10**. Overall assignment accuracies to the two main *T. h. hermanni* groups, Italian Peninsula (ITP, NCA and CCA) and Mediterranean Islands (SCA, SIC and SAR), estimated via Monte Carlo cross-validation of the 96 SNPs panels, with two levels of training (baseline) individuals (50% and 70% of individuals from each group) crossed by up to three levels of training loci (top 50%, 75% and all loci) by 500 resampling events: (a) PCA; (b) $F_{ST}$; and (c) RF.
