## Supplementary Methods for "From STRs to SNPs via ddRAD-seq: geographic assignment of confiscated tortoises at reduced costs"

### ddRAD library preparation and Ion Torrent PGM sequencing

*In silico* digestions with the R package simRAD (Lepais and Weir, 2014) were performed on the available genome of *Chrysemys picta* (Badenhorst et al., 2015) to evaluate the best combination of restriction enzymes for the ddRAD-seq experiment. The simulations suggested the pair *Sbf*I and *Nco*I as an optimal combination to obtain about 5000 fragments around 300 ±50 bp.

We prepared five libraries of 10-22 samples each, for 5 Ion Torrent PGM chips. Firstly, 500 ng of DNA for each sample were digested at 37° C for 3 h in a 30 μl reaction volume containing 10 U of *Sbf*I-HF (New England Biolabs), 10 U of *Nco*I-HF and CutSmart Buffer 1X. Digested samples were then purified with 1.5X Agencourt AMPure XP reagent (Beckman Coulter) following manufacturer's instructions.

Each purified sample was ligated with barcoded Adapter A and Y-shaped Adapter P1 (both modified for sticky-end ligation with *SbfI* and *NcoI* cut, respectively) in a 40 μl reaction volume containing 300 ng of purified digested DNA, 400 U T4 DNA Ligase (New England Biolabs), 1X T4 Buffer and 2 μl of working solution of Adapter A and Adapter P1. The working solutions of adapters were calculated as in (Peterson et al., 2012). Ligation reaction was performed in a thermalcycler with the following steps: 23° C for 30 min, 65° C for 10 min and 41 cycles of 45 sec each starting at 64° C with a decrement of 1° C at each cycle. After ligation, all samples of each library were pooled and purified with 1.2X Agencourt AMPure XP reagent (Beckman Coulter) following manufacturer's instructions.

We performed size selection of 300 bp library fragments on a E-gel Size Select 2% Agarose Gel (Invitrogen). Size selected fragments were PCR amplified in a reaction containing Phusion HF Polymerase (2 U/μl, New England Biolabs) 0.2 μl, 5X Phusion HF Buffer 5 μl, dNTP (10 μM) 0.5 μl, Ion Torrent Primer A (10 μM) 2 μl, Ion Torrent Primer P1 (10 μM) 2 μl and size selected library to a final volume of 25 μl. Amplification conditions were the following: initial denaturation at 98° C for 1 min, 12 cycles of 10 sec at 98° C, 20 sec at 62° C and 1 min at 72° C and a final extension of 5 min at 72° C. We amplified 4 replicates for each library that were pooled and purified with 1.5X Agencourt AMPure XP reagent (Beckman Coulter).

Each library was quantified with Qubit using the dsDNA BR kit (Invitrogen) and diluted to 26 pM for the Ion Torrent PGM sequencing workflow.

Libraries were then amplified in emulsion PCR on a One Touch 2 System with the Ion PGM Hi-Q OT2 kit (Life Technologies) and enriched with the Ion One Touch ES following manufacturer's instructions. The sequencing was performed on the Ion Torrent PGM with the Ion PGM Hi-Q View Sequencing kit (Life Technologies). Two libraries were sequenced on Ion 316 Chips and 3 on Ion 318 Chips.
